# Intrinsic immunity against HAdV is achieved by a novel epigenetic silencing complex

**DOI:** 10.1101/2025.02.10.637372

**Authors:** Julia Mai, Maryam Karimi, Anna Kuderna, Anna Heider, Thomas Günther, Caroline C. Friedel, Adam Grundhoff, Thomas Stamminger, Sabrina Schreiner

**Author notes:** all authors contributed equally to this work (Julia Mai, Maryam Karimi, Anna Kuderna).

## Abstract

The DNA double-stranded genome of human adenoviruses (HAdV) is preferentially targeted by host factors involved in chromatin remodeling, such as SPOC1 and KAP1 to inhibit efficient viral gene expression. HAdV genomes undergo alterations through association with host histones and epigenetic modifications; however, the precise underlying mechanism remains elusive. A recently discovered silencing mechanism for retrotransposons and retroviruses involves the Human Silencing Hub (HUSH) complex. This complex of MPP8, TASOR, and PPHLN1 safeguards the human genome by utilizing histone H3 Lys9 trimethylation (H3K9me3) to block transcription. Through the recruitment of SETDB1 and MORC2, the HUSH complex silences host genes and condenses target genomes to combat infections such as HIV, MLV, and AAV. Here, we present evidence that the HUSH complex effectively restricts HAdV infection. To counteract the repressive function of this epigenetic silencing complex, HUSH factors are inhibited through binding of HAdV proteins and subsequent relocalization. We observe that MPP8 is targeted by the adenoviral E3 ubiquitin ligase, thus recruited by the viral early proteins E1B-55K and E4orf6 for proteasomal degradation. In summary, we provide evidence that the HUSH complex is a previously unrecognized host factor that restricts HAdV gene expression and replication. Based on these novel findings, we propose that HUSH represents a promising therapeutic target to combat HAdV infection.

## Introduction

The human silencing hub complex (HUSH) has been identified as a multiprotein complex that regulates epigenetic silencing of genes in heterochromatic regions [1]. This complex consists of the transcriptional activation suppressor (TASOR), M-phase phosphoprotein 8 (MPP8) and periphilin (PPHLN1), which are attracted to regions of the genome with high levels of H3K9me3 [1] and promote an increase in H3K9me3 by interaction with SET domain bifurcated 1 (SETDB)1 and the CW zinc finger protein 2 (MORC2) of the MORC family. SETDB1 is a histone methyltransferase that deposits H3K9me3 at specific sites, and MORC2 is an ATP-dependent chromatin remodeling protein that compacts chromatin [1, 2]. However, the function and regulation of this complex are not yet sufficiently understood. Previous publications suggest that the HUSH complex is recruited to the specific sites in two different ways: RNA- mediated and DNA-mediated HUSH recruitment [3, 4]. The first is PPHLN1-dependent RNA- induced silencing, in which reverse transcription products are used to achieve silencing. The second, non-canonical method relies on the activities of the chromodomains of nuclear protein 220 (NP220) and MPP8, which bind to the non-integrated DNA of MLV (murine leukaemia virus) and AAV (adeno-associated virus) DNA. This binding triggers the recruitment of other factors that work together to silence the target loci [4].

The HUSH subunit MPP8 is a mitotic phosphoprotein containing an amino-terminal chromodomain and a carboxy-terminal ankyrin domain [5]. It is crucial for the organization of heterochromatin as it is preferentially localized in the regions mediating the interaction between de novo DNA methylation and histone H3K9 methylation [6]. The MPP8 chromodomain interacts with GLP in a methylation-dependent manner, and by forming a dimer, it may link methylated Dnmt3a and GLP to create a repressing complex that forms heterochromatin [5]. In the context of viral infections, the HUSH complex has been identified as an antiviral factor that regulates endogenous and viral genes of HIV and MLV [1, 7, 8]. To counteract these functions, the viral proteins Vpx and Vpr act as adaptors for the E3 ubiquitin ligase DCAF1- CUL4A/B to reduce the protein concentrations of the HUSH components [8–11]. Recruitment of the HUSH complex to non-integrated viral DNA is facilitated by NP220, which has been identified as a key protein in this process [12]. NP220 is a nuclear protein with a specific affinity for cytidine clusters on double-stranded DNA. It consists of a DNA-binding domain (DB) and a C-terminal zinc finger motif of type C2H2 (ZnF) [13]. The exact role and contribution of the HUSH complex in relation to non-integrated DNA viruses, such as human adenoviruses (HAdV), is still unclear and the subject of further research. However, in a recent study, the HUSH complex was identified as a potential factor in the regulation of latent HSV-1 infection [14]. During HSV-1 infection, promyelocytic leukemia bodies (PML-NBs) recruit viral genomes upon viral entry and form viral DNA-containing NBs (vDCP-NBs) [15, 16]. These vDCP-NBs have been shown to be characterized by repressive chromatin that is dependent on the HUSH complex, as depletion of these components prior to viral infection leads to a decrease in latent HSV-1 genomes, whereas overexpression restricts viral gene expression [14, 16].

HAdV are ds-DNA viruses that can cause sporadic epidemics and, depending on the virus type and corresponding tropism, can cause a variety of different diseases [17–20]. In recent years, highly pathogenic types have frequently led to severe courses of disease in immunocompetent patients, but no specific therapy or immunization is yet available to the general population [21, 22]. Since the regulation of viral gene expression is still a scientific black box, especially with regard to newly evolving HAdV types it is of great interest to identify host factors that shed light on how HAdVs control their transcription in order to develop new therapeutic. Previous studies have already shown that active gene expression can be inhibited by restrictive host factors [23–27]. Human adenoviruses have therefore developed a mechanism with which they can inhibit these factors. The viral proteins E1B-55K and E4orf6 cooperate with the cellular factors Elongin B/C, Rbx1 and Cullin5 to form an E3 ubiquitin ligase that marks the negative host proteins for proteasomal degradation [24, 28–34]. This is crucial for efficient virus replication, but not all aspects of this regulation are yet known. Here, we investigated the role of the HUSH complex, a novel heterochromatin regulator, on HAdV replication and gene expression. In summary, the study highlights how the HUSH complex, particularly MPP8, regulates HAdV transcription and is counteracted by viral proteins like E1B-55K, which degrade HUSH components to facilitate efficient viral gene expression and replication.

In summary, this study highlights how the HUSH complex, particularly MPP8, regulates HAdV transcription and is counteracted by viral proteins like E1B-55K, which degrade HUSH components to facilitate efficient viral gene expression and replication.

## Results

### Protein levels of MPP8 are decreased during HAdV infection

The HUSH complex serves as an important epigenetic regulator in viral infections, silencing viral elements through histone modifications [14, 35]. Modulation of its protein content is closely related to the stability of its core components and their interactions, which can influence the ability of the complex to repress viral transcription [8, 36]. To investigate the regulation of the HUSH complex during HadV infection, we first analyzed the gene expression of NP220 and HUSH components MPP8, TASOR, and PPHLN1 in western blot analysis. Here, we infected human lung carcinoma H1299 cells with HadV-wt and harvested at the indicated time points during the course of infection (Fig. 1A). The early viral proteins E1B-55K and E2A serve as infection control and are detectable 24 h post-infection (pi) with an increase in protein levels during later time points of HadV replication (Fig. 1A lane 4-6). Our results show a strong increase in NP220 protein levels during HadV infection (Fig. 1A, compare lanes 4-6 to lanes 1-3). NP220 levels are even more elevated during the course of infection peaking at 48 h and 72 h pi (Fig. 1A, lanes 4-6). On the other hand, we observed reduction of protein levels of all three HUSH components at the latest 48 h pi. MPP8 expression is severely impaired at 48 and 72 h pi compared to the corresponding uninfected control samples with MPP8 expression not detectable at 72 h pi (Fig. 1A, compare lane 5 to lane 2, compare lane 6 to lane 3). Inhibition of TASOR and PPHLN1 were less pronounced, however, significantly reduced 48 h and 72 h pi compared to endogenous steady state levels in uninfected cells (Fig. 1A, compare lane 5 to lane 2, compare lane 6 to lane 3).

**Fig. 1:**
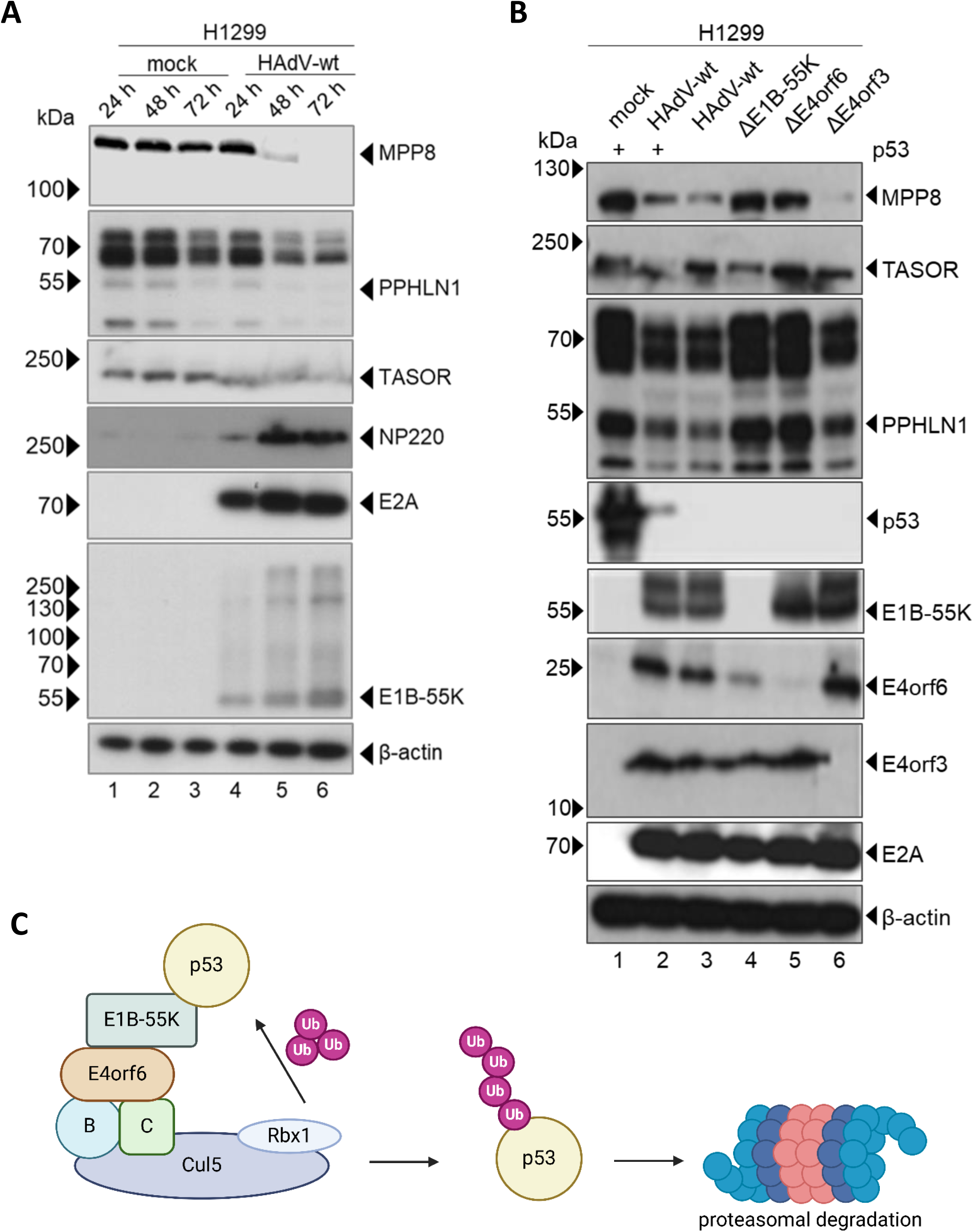
Protein levels of HUSH components are declined during HAdV infection. **(A)** H1299 cells were infected with HAdV-wt at a MOI of 50 and harvested 24 h, 48 h and 72 h pi. Whole cell lysates were prepared, separated by SDS PAGE, and subjected to immunoblotting using mAb AC-15 (β-actin), rabbit pAb MPP8, rabbit pAb PPHLN1, rabbit pAb TASOR, rabbit pAb NP220, mouse mAb E2A, and mouse mAb E1B-55K. β-actin serves as a loading control. Molecular weights in kDa are indicated in the left, proteins are indicated on the right. **(B)** H1299 cells were infected with HAdV-wt and mutant versions depleted for early viral proteins, ΔE1B- 55K, ΔE4orf6, and ΔE4orf3, at a MOI of 50. In addition, we transiently transfected two samples with p53 DNA as indicated. Cells were harvested 48 h pi and whole cell lysates were prepared. Proteins were separated by SDS PAGE and subjected to immunoblotting using mAb AC-15 (β-actin), mouse mAb MPP8, rabbit pAb PPHLN1, rabbit pAb TASOR, mouse mAb E2A, mouse mAb E1B-55K, mouse mAb E4orf6, and rat mAb E4orf3. β-actin serves as a loading control. Molecular weights in kDa are indicated in the left, proteins are indicated on the right. **(C)** Illustration of the viral E3 ubiquitin ligase targeting p53 for proteasomal degradation during HAdV infection. Created with *Biorender.com*.

Since we have demonstrated a reduction of protein levels by HadV, we were interested which viral factors modulate protein expression levels of MPP8, TASOR and PPHLN1 (Fig. 1A). Therefore, we infected H1299 cells with HAdV-wt and mutant variants lacking the expression of early viral proteins E1B-55K, E4orf6 or E4orf3 (Fig. 1B). These are known factors involved in the suppression of restrictive host factors [23, 31, 37–39]. As an additional control for a host factor degraded by E1B-55K and E4orf6 [33] (Fig. 1C), we have transiently transfected p53 into the p53-negative H1299 cells. We observed E1B-55K/E4orf6 mediated proteasomal degradation of p53 during HadV-wt infection as expected and consistent with published data [33] (Fig. 1B, compare lane 1 to lane 2, Fig. 1C). As infection control and to characterize the virus mutant variants, we analyzed protein levels of E2A, E1B-55K, E4orf6 and E4orf3. The corresponding depleted gene products were not expressed by the viruses used and E2A indicated comparable virus concentrations in our experiment (Fig. 1B). As expected, we observed a reduction in protein levels of MPP8, TASOR and PPHLN1 during HAdV-wt infection (Fig. 1B, compare lane 3 to lane 1), which is independent of p53 expression (Fig. 1B, compare lane 3 to lane 1). We find that the MPP8, TASOR and PPHLN1 protein concentrations are also reduced in the E4orf3-depleted virus (ΔE4orf3) compared to HAdV-wt infection (Fig. 1B, compare lane 6 to lane 3 and 1). However, we observe that during ΔE1B-55K (Fig. 1B, compare lane 4 to lane 3 and 1) and ΔE4orf6 (Fig. 1B, compare lane 5 to lane 3 and 1) infection, all HUSH regulator proteins proteins MPP8, TASOR and PPHLN1 do not show reduced levels but comparable protein levels than observed in uninfected samples. In sum, we report here that levels of the HUSH components MPP8, TASOR and PPHLN1 are reduced during HAdV infection, being independent of p53 and E4orf3 expression but mediated by E1B- 55K and E4orf6 expression.

### MPP8 is targeted by the viral E3 ubiquitin ligase for proteasomal degradation

Next, we examined the downregulation of HUSH components MPP8, TASOR, and PPHLN1 by HAdV. The early viral proteins E1B-55K and E4orf6 build an E3 ubiquitin ligase with cellular proteins, which can label proteins for proteasomal degradation [31–33, 38–42]. Since we observe no HUSH modulation in cells infected with mutant virus variants that lack E1B-55K or E4orf6 protein expression, we hypothesized that the respective host regulators are degraded by the viral E3 ubiquitin ligase and degraded proteasomally (Fig. 1B, C). We infected H1299 cells with HAdV-wt and treated these cells with MG132 proteasome inhibitor 16 h prior to harvesting. After preparation of whole-cell lysate we subjected the samples to immunoblotting. E2A serves as an infection control being expressed in HAdV-infected cells independent of the MG132 treatment (Fig. 2A, compare lane 2 and lane 4). To verify inhibition of the proteasome by MG132, we analyzed ubiquitin protein levels in our samples. Cells treated with MG132 show a stronger expression of upcoming ubiquitin bands in our immunoblots, as the ubiquitinylated proteins can no longer be degraded by the proteasome (Fig. 2A, compare lanes 3 and 4 to lane 1 and 2). Notably, we observed different effects on HUSH protein levels after proteasome inhibition. PPHLN1 showed no difference between treated and untreated cells (Fig 2A, compare lane 2 and lane 4). In both samples, protein levels were reduced in HAdV-infected cells compared to the control (Fig. 2A, compare lane 2 and 4 to 1). Both TASOR and MPP8 could be identified as target proteins for the host proteasome during infection (Fig. 2A, compare lane 4 to lane 2). Both proteins show elevated protein levels in HAdV-infected cells after MG132 treatment compared to the untreated sample. This can be obtained most clearly for MPP8 protein expression, which was no longer detectable in the infected, untreated sample, while the protein concentration increased again after suppression of the host proteasome mediated degradation (Fig. 2A, compare lane 4 to lane 2).

**Fig. 2:**
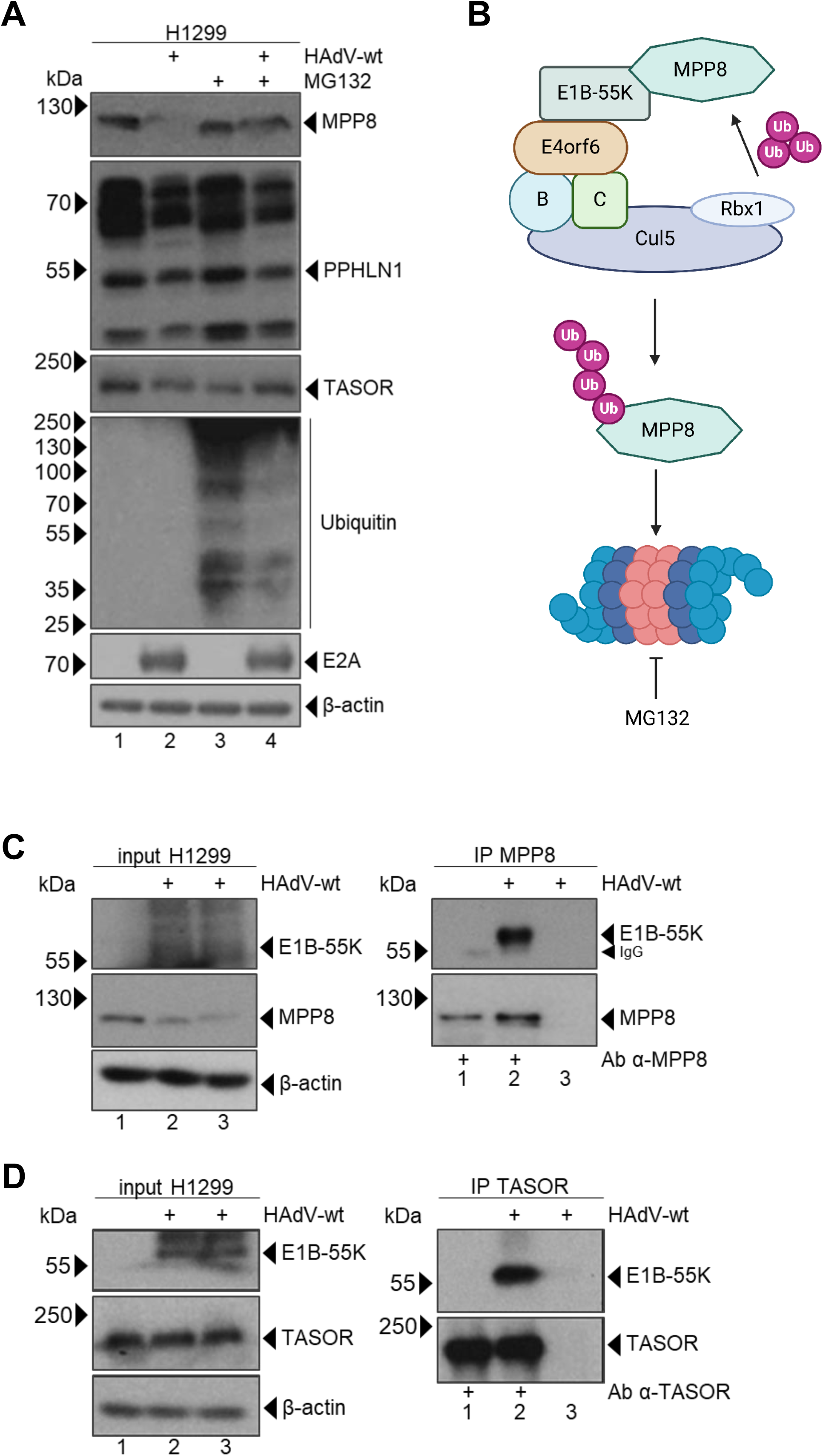
Proteins of the HUSH complex are a target for viral E3 ubiquitin ligase. **(A)** H1299 cells were infected with HAdV-wt at a MOI of 50 and treated with proteasome inhibitor MG132. Cells were harvested 48 h pi and whole cell lysates were prepared. Proteins were separated by SDS PAGE and subjected to immunoblotting using mAb AC-15 (β-actin), mouse mAb MPP8, rabbit pAb PPHLN1, rabbit pAb TASOR, mouse mAb Ubiquitin and mouse mAb E2A. β-actin serves as a loading control. Molecular weights in kDa are indicated in the left, proteins are indicated on the right. **(B)** Graphical representation of the viral E3 ubiquitin ligase composed of early viral proteins E1B-55K and E4orf6. MPP8 is targeted by E1B-55K and E4orf6 for proteasomal degradation, which can be inhibited by MG132 treatment. **(C, D)** H1299 cells were infected with HAdV-wt at a MOI of 50 and harvested 48 h pi. Co-immunoprecipitation assay was performed by using a **(C)** mouse mAb MPP8 or **(D)** rabbit pAb TASOR. Proteins were separated by SDS PAGE and subjected to immunoblotting using mAb AC-15 (β-actin), mouse mAb E1B-55K, **(C)** mouse mAb MPP8 or **(D)** rabbit pAb TASOR. β-actin serves as a loading control. Molecular weights in kDa are indicated in the left, proteins are indicated on the right.

To target host restriction factors, viral E1B-55K and E4orf6 cooperate with host proteins Cullin5, Elongin B/C and Rbx1. E4orf6 bind via its BC box motif to the Elongins B/C and interacts with E1B-55K, which recruits the target proteins (Fig. 2B) [32, 41, 42]. For Daxx, we recently reported that E1B-55K alone is sufficient to assemble an E3 ubiquitin ligase to ubiquitinylate the host target [31]. Next, we, we performed co-immunoprecipitation analyses to gain insight into E1B-55K association with HUSH factors. We precipitated MPP8 (Fig. 2C) and TASOR (Fig. 2D) from HAdV-infected H1299 cells using specific antibodies. Our input samples demonstrate successful infection of the cells with HAdV-wt by detecting E1B-55K (Fig. 2C, D, input lane 2 and 3). The samples derived from the immunoprecipitation were stained for the pulled protein to show pulldown of the target protein (Fig. 2C, D, IP lane 2). In addition, we controlled immunoprecipitation without adding specific antibodies to exclude non-specific binding of E1B-55K (Fig. 2C, D, IP lane 3). We detected E1B-55K signals in our immunoprecipitations indicating an interaction of MPP8 and TASOR with the E3 ubiquitin ligase proteins (Fig. 2C, D, lane 2). In summary, we provide evidence that PPHLN1 is reduced during infection via a proteasome-independent mechanism (Fig.2A). MPP8 and TASOR both bind to E1B-55K and undergo proteasomal degradation (Fig. 2A, B, C).

### Depletion of MPP8 protein expression promotes HAdV replication

During HAdV infection, host proteins are proteasomally degraded by the E1B-55K/E4orf6 E3 ubiquitin ligase complex to suppress host antagonistic function and promote virus gene expression [31, 33, 42]. Consequently, we next investigated the effect of the HUSH complex on HAdV replication. We have generated knockdown cell lines for MPP8, TASOR or PPHLN1. These cells were infected with HAdV-wt and harvested 48 h pi prior to collection of whole cell lysates to purify viral progeny. Virus yield analysis showed that neither knockdown of TASOR nor PPHLN1 had any effect on virus replication. However, we observed a significant increase in newly synthesized virions after MPP8 depletion (Fig. 3A). MPP8 (Fig. 3B, compare lane 3 and 4 to lane 1 and 2), PPHLN1 (Fig. 3B, compare lane 5 and 6 to lane 1 and 2), and TASOR (Fig. 3B, compare lane 7 and 8 to lane 1 and 2) expression as well as HAdV infection was controlled by immunoblotting (Fig. 3B). As we observed the most significant effect on HAdV progeny in MPP8 depleted cells, we further investigated RNA levels in these cells. We observed a 1.75-fold increase in Hexon mRNA (Fig. 3D), while E1A transcript expression was significantly increased by 1.5 upon in our qPCR experiments in MPP8 depleted cells (Fig. 3C). To investigate the effect of MPP8 expression on viral proteins we immunostained early proteins E1A, E1B-55K, E2A, E4orf6, and E4orf3, and late viral capsid proteins (Fig. 3E). We could not detect significant changes for E1A, E2A, E4orf6, and E4orf3, however, E1B-55K and capsid were significantly increased in infected shMPP8 compared to shCRL cells (Fig. 3E, compare lane 4 to lane 2; Fig. 3F).

**Fig. 3:**
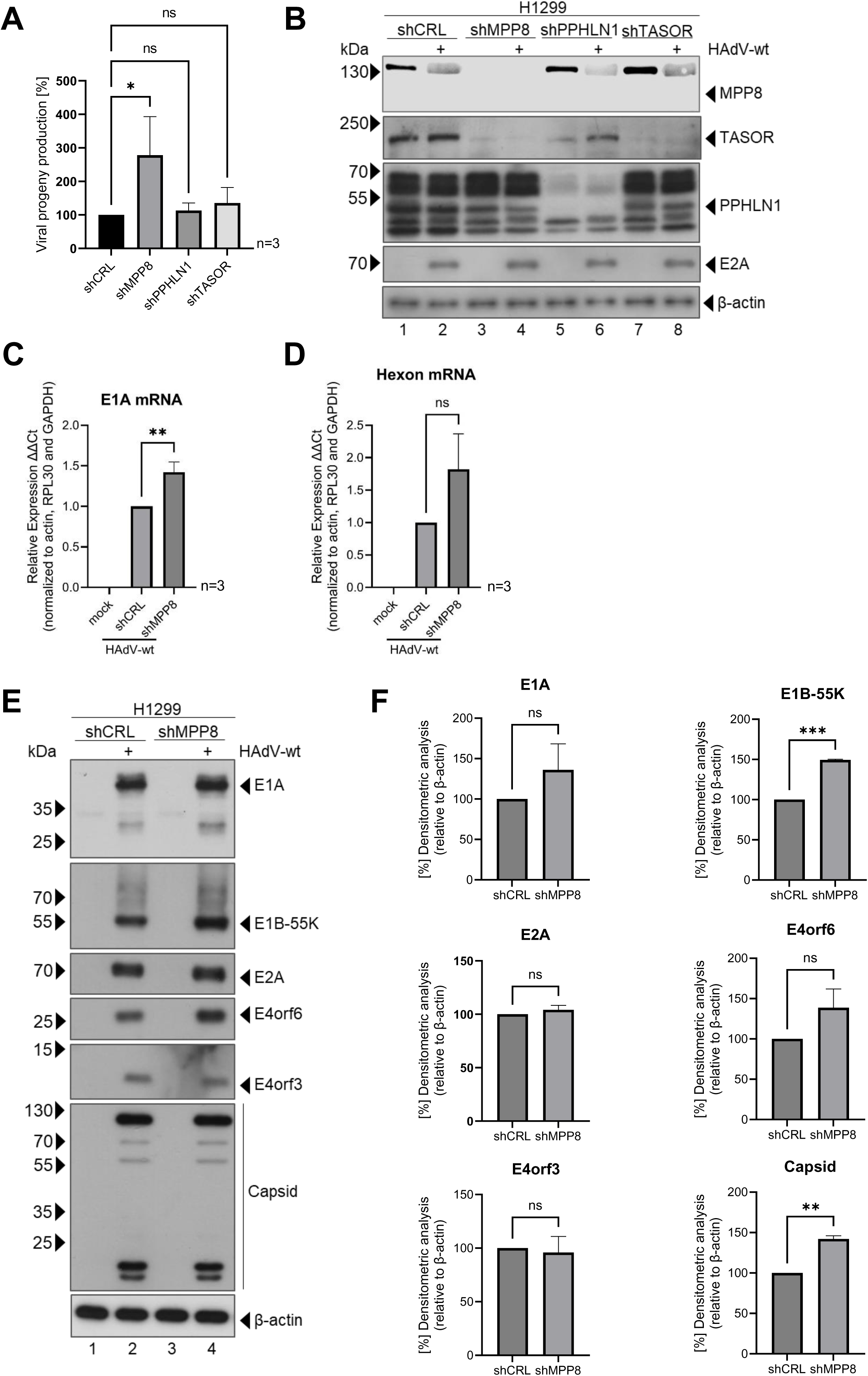
HAdV infection and replication is promoted by knockdown of MPP8. H1299 shCRL, shMPP8, sh PPHLN1, and shTASOR were infected with HAdV-wt at a MOI of 50 and harvested 48 h pi. **(A)** Viral particles were isolated from HAdV-infected cells by three cycles of freeze and thaw. HEK293 cells were reinfected with a serial dilution of the viral particles and virus yield was determined using quantitative E2A immunofluorescence staining. Bar charts represents mean values of three independent biological replicates. H1299 shCRL cells were set to 100% and the remaining samples were normalized. Statistically significant differences were determined using an one-way ANOVA and Dunnet’s T3 test with the GraphPad Prism10 software. ns = not significant; \**p* ≤ 0.05. **(B)** Whole cell lysates were prepared from some of the cells, from which virus particles were isolated. Proteins were separated by SDS PAGE and subjected to immunoblotting using mAb AC-15 (β-actin), rabbit pAb MPP8, rabbit pAb PPHLN1, rabbit pAb TASOR, and mouse mAb E2A. β-actin serves as a loading control. Molecular weights in kDa are indicated in the left, proteins are indicated on the right. **(C,D)** Total mRNA was isolated using TRIzol reagent, reverse transcribed and subjected to qPCR using oligonucleotides specific for viral transcripts **(C)** E1A and **(D)** Hexon. The data was normalized on the respective GAPDH, actin and RPL30 mRNA levels. Bar charts represents mean values of three independent biological replicates measured in technical triplicates. Statistically significant differences were determined using an unpaired student’s t test with the GraphPad Prism10 software. ns = not significant; \*\**p* ≤ 0.01. **(E)** Whole cell lysates were prepared, proteins separated by SDS PAGE and subjected to immunoblotting using mAb AC- 15 (β-actin), rabbit pAb MPP8, mouse mAb E1A, mouse mAb E1B-55K, mouse mAb E2A, mouse mAb E4orf6, rat mAb E4orf3, and rabbit pAb Capsid. β-actin serves as a loading control. Molecular weights in kDa are indicated in the left, proteins are indicated on the right. **(F)** Densitometric analysis of detected band was performed using *ImageJ* (version 1.54m) to quantify protein levels. Relative protein levels were normalized to the respective housekeeping β-actin steady state levels. Bar charts represent mean values and standard deviations are based on two independent experiments. Statistically significant differences were determined using an unpaired student’s t test with the GraphPad Prism10 software. ns = not significant; \*\**p* ≤ 0.01; \*\*\**p* ≤ 0.001.

### HAdV infection recruits MPP8 to PML tracks and HAdV replication centers

Recent studies have identified the HUSH complex as a factor that regulates vDCP-NBs and thus viral gene expression and latency of HSV-1 [14, 43]. Similar to HSV-1, HAdV also localize their DNA genome juxtaposed to PML-NBs, an event that is essential for viral replication [44, 45]. Since MPP8 is a factor associated with DNA to mediate H3K9 methylation, we investigated the localization of MPP8 during HAdV infection (Fig. 3). We infected H1299 cells with HAdV- wt, fixed the cells and triple-stained for E4orf3, E2A and MPP8. E4orf3 is an early viral protein known to interact and to relocalize PML-NBs into track-like structures [44], while the DNA-binding protein E2A is the viral DNA binding protein and a marker for adenoviral replications centers (RCs) [45]. Both viral proteins were not detectable in uninfected cells (Fig. 4 A, panels a-t). As expected, 24 h pi we observe E4orf3-containing PML tracks (Fig. 4A, panels v, aa, af, ak, and ap; Fig. 4B, panels a and d) and dot- / ring-like E2A RCs (Fig. 4A, panels w, ab, ag, ai, and ao; Fig. 4B, panels b and e). MPP8 localizes to the nucleus as numerous small dots in uninfected cells (Fig. 4A, panels d, i, n, and s). Intriguingly, MPP8 localization was altered in HAdV-infected cells. We observe that MPP8 is recruited to E4orf3-containing PML tracks (indicated by white arrows; Fig. 4A, panels x, ac, ad, ah, ai, am, an and ap; Fig. 4B, panels c and f) and also to adenoviral RCs (indicated by yellow arrows; Fig. 4A, panels x, ac, ad, ah, am and ao; Fig. 4B, panels c and f).

**Fig. 4:**
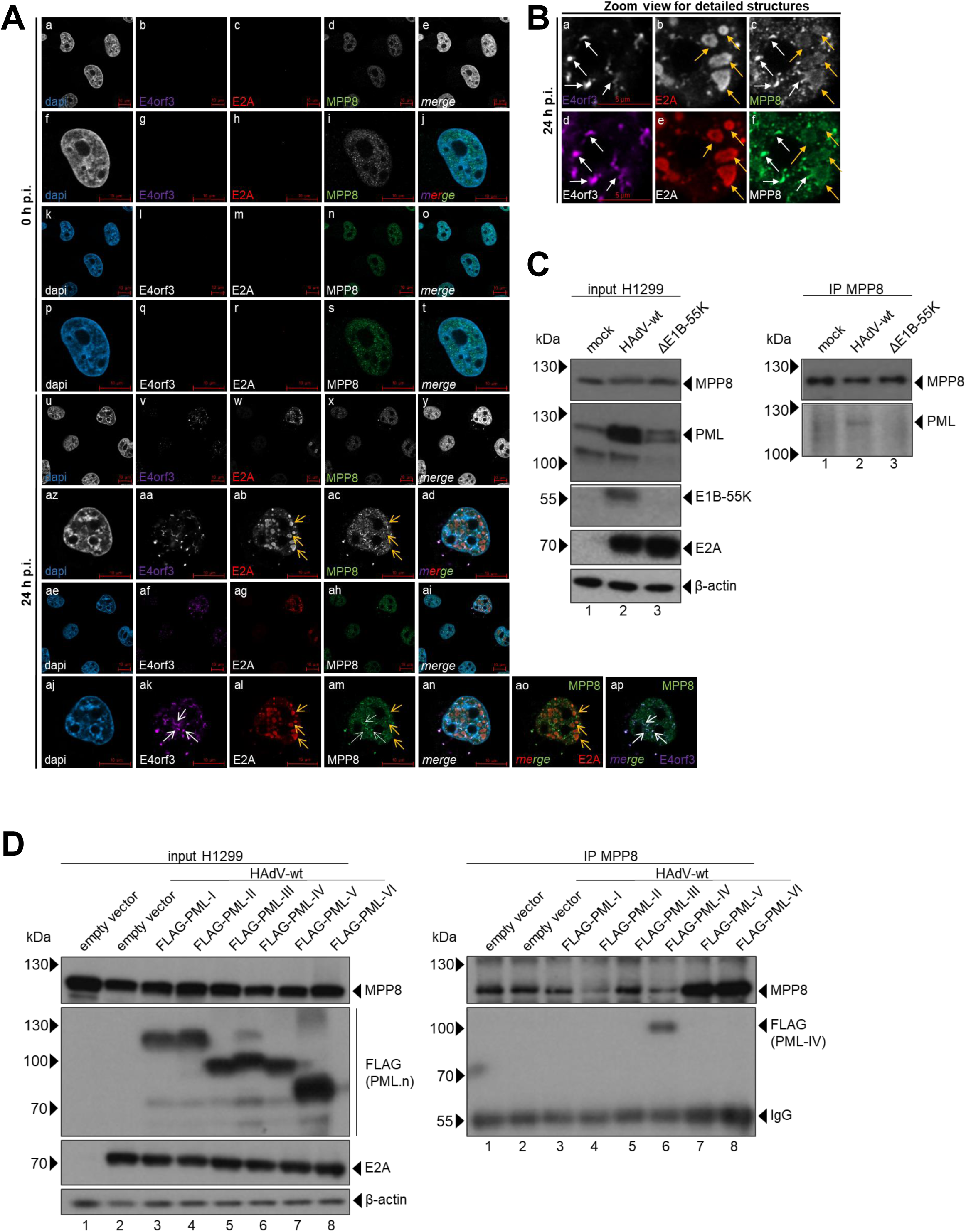
MPP8 is localizing to PML tracks and adenoviral replication centers (RCs). **(A, B)** H1299 cells were infected with HAdV-wt at a MOI of 50 and fixed with 4% paraformaldehyde 0 h and 24 h pi. Cells were permeabilized with 0.5% triton X100 and blocked with TBS-BG. Cells were triplestained using rat pAb E4orf3, rabbit pAb E2A, and mouse mAb MPP8. Primary antibodies were detected with Alexa647- (magenta; α-E4orf4), Alexa568- (red; α-E2A), and Alexa488- (green; α-MPP8) conjugated secondary antibodies. Images were taken with a *Zeiss* LSM 900 Airyscan 2. **(A)** Cells are displayed in two different ways: grayscale (a-j, u-ad) and colored (k-t, ae-ap). Ovierview images are shown in a-e, k-o, u-y, and ae-ai. Zoom view of single cells is shown in f-j, p-t, az-ad, and aj-ap. Merge images of all three stainings is shown in j, o, t, ad, ai, and an. Panel ao shows merge of E2A and MPP8, panel ap shows merge of E4orf3 and MPP8. Red scale bar represents 10 µm. White arrows mark E4orf3-containing PML tracks localizing with MPP8. Yellow arrows mark E2A-marked RCs localizing with MPP8. **(B)** Images show a detailed zoom of the cell in (A) panels aj-ap. The cell is displayed in two different ways: grayscale (a-c) and colored (d-f). Red scale bar represents 5 µm. White arrows mark E4orf3-containing PML tracks localizing with MPP8. Yellow arrows mark E2A-marked RCs localizing with MPP8. **(C)** H1299 cells were infected with HAdV-wt or HAdV ΔE1B-55K at a MOI of 50 and harvested 48 h pi. Co-immunoprecipitation assay was performed by using a mouse mAb MPP8. Proteins were separated by SDS PAGE and subjected to immunoblotting using mAb AC-15 (β-actin), mouse mAb E1B-55K, mouse mAb E2A, mouse mAb MPP8, and rabbit pAb PML. β-actin serves as a loading control. Molecular weights in kDa are indicated in the left, proteins are indicated on the right. **(D)** H1299 cells were infected with HAdV-wt at a MOI of 50 and transiently transfected with an empty vector control or FLAG-tagged PML isoforms (PML.n). The samples were harvested 48 h pi and co-immunoprecipitation assay was performed by using a mouse mAb MPP8. Proteins were separated by SDS PAGE and subjected to immunoblotting using mAb AC-15 (β-actin), mouse mAb E2A, mouse mAb MPP8, and rabbit pAb FLAG. β-actin serves as a loading control. Molecular weights in kDa are indicated in the left, proteins are indicated on the right.

### HAdV E1B-55K mediated binding of MPP8 to PML during infection

To further verify these immunofluorescence studies, we performed co-immunoprecipitation assays in HAdV-wt and HAdV ΔE1B-55K-infected H1299 cells (Fig. 4C). Comparable infection with both viruses was verified by E2A staining, while E1B-55K was not expressed by the depletion variant (Fig. 4C, input compare lane 2 with lane 3). Pulldown of protein complexes was achieved by a MPP8 antibody. We obtained MPP8 staining in all samples (Fig. 4C), and efficient direct immunoprecipitation with MPP8 was detected (Fig. 4C, IP MPP8). Subsequent staining of PML then showed an interaction of MPP8 only in HAdV-wt-infected cells, but not in uninfected cells (Fig. 4C, IP MPP8 compare lane 1 to lane 2) or in cells infected with the E1B- 55K depleted virus variant (Fig. 4C, IP MPP8 compare lane 3 to lane 2). E1B-55K is a multifunctional factor that interacts with PML isoforms IV and V, amongst others, which is crucial for the inactivation of p53 [46].

### MPP8 is recruited by PML-IV isoform and complexes with E1B-55K

To further decipher the association of MPP8 with PML and the interplay with E1B-55K, we performed co-immunoprecipitation analyses in which we further examined the interaction of MPP8 with the different PML isoforms. Thus, we infected cells with HAdV-wt and transiently transfected them with single FLAG-tagged PML isoforms. To precipitate the protein complexes we used an antibody targeting the MPP8 protein (Fig. 4 D). Expression of all investigated proteins and HAdV infection was verified in our input samples (Fig. 4D, input H1299). After successful pulldown of the MPP8 protein and subsequent staining for the FLAG-tag of the single PML isoforms, we confirm an interaction of MPP8 with PML-IV (Fig. 4D, IP MPP8 lane 6). In summary, MPP8 is relocalized to adenoviral RCs and associates with PML-IV in tracks dependent on E1B-55K.

## Discussion

This is the first study to show a role of the HUSH complex on HAdV infection. Particularly in the context of the heterochromatin component MPP8 our study offers valuable insights into the regulation of viral gene expression (Fig. 3). Our findings suggest that knockdown of MPP8 promotes viral replication (Fig. 3A) and viral transcription (Fig. 3C-F). Since MPP8 mediates H3K9 trimethylation [47–49] in cooperation with the HUSH complex, we hypothesize that the HUSH complex promotes heterochromatin formation of the adenoviral genome. This complex is highly regulated and its functionality depends on the expression of the individual subunits and interaction partners [13]. TASOR protein levels were strongly dependent on MPP8 expression (Fig. 3B), suggesting a possible interplay between MPP8 and TASOR to regulate HAdV replication. Conversely, the absence of TASOR alone does not affect efficient adenoviral infection (Fig. 3A), since we observed similar MPP8 levels and efficient viral replication as in shCRL cells (Fig. 3A, B). However, this does not exclude an important role of TASOR in virus infection. To counteract any restriction by HUSH, HAdV targets the components of the HUSH complex to suppress antiviral capacity (Fig. 1, Fig. 2). In summary, our study provides evidence for a role of MPP8 on adenoviral transcription, most presumably even in connection with TASOR.

Our immunoprecipitation assays revealed an association of MPP8 and TASOR with the viral E1B-55K. This early viral protein assembles together with E4orf6 and additional cellular host factors an E3 ubiquitin ligase marking target proteins for proteasomal degradation [31–33, 41, 42]. In our infection studies, we found that both viral proteins are required for the degradation of MPP8 and TASOR, while other viral proteins have no effect (Fig. 1B). These data provide compelling support for the degradation of HUSH proteins after ubiquitinylation by the viral E3 ubiquitin ligase. This is further supported by the stabilization of MPP8 and TASOR after inhibition of the proteasome (Fig. 2A). In contrast, we did not observe any involvement of the proteasome on PPHLN1 protein levels (Fig. 2A). However, a role of the two proteins E1B-55K and E4orf6 can also not be excluded for PPHLN1 (Fig. 1B). First, the expression of PPHLN1 could already be reduced at the transcriptional level. Furthermore, degradation at the translational level does not have to occur via the proteasome. This can also occur via lysosomal, caspase-mediated, calpain-mediated or mitochondrial degradation [50–52]. E1B- 55K is not only a part of the E3 ubiquitin ligase, but can also function as an E3 SUMO ligase that modulates the post-translational modification (PTMs) of proteins. Therefore, modification of PPHLN1, such as phosphorylation or SUMOylation might alter protein stability and localization, as we have previously published for other host target proteins such as KAP1, Daxx and p53 [30, 39, 46, 53]. Since our lysate preparation limits the detection of insoluble proteins, future experiments, such as qPCR, immunofluorescence, fractionation assays, and detection of PTMs, will provide further insights on the regulation of PPHLN1 by HAdVs. In addition, the reduction of PPHLN1 could also be a secondary effect through the modulation of other host factors such as transcription factors by E1B-55K and E4orf6, however, the gene expression of PPHLN1 is poorly understood to date. In conclusion, our findings suggest proteasomal degradation of MPP8 and TASOR via the viral E3 ubiquitin ligase to promote efficient viral replication and gene expression.

While PPHLN1 cannot bind MPP8, TASOR associates with PPHLN1 and MMP8 and forms the HUSH complex. MPP8 can then interact with NP220, which binds directly to linear unintegrated DNA. Previous studies have shown that the association of the HUSH complex is reduced after silencing of MPP8. HUSH components TASOR and PPHLN1 are no longer associated with the DNA and methylation of H3K9me3 is diminished [13]. This might be an explanation for how HAdV gene expression is promoted after MPP8 knockdown (Fig. 3) and for the underlying mechanism how HAdV represses HUSH complexes, since we observed destabilization of the MPP8 protein (Fig. 1A). However, limitations in the significant induction of gene expression for certain viral transcripts (Fig. 3D) and viral proteins (Fig. 3E, F) could be due to NP220, which was shown to be induced during HAdV infection (Fig. 1A). Since NP220 is the most important protein that binds DNA, gene expression could be suppressed by the interaction of NP220 with certain histone deacetylases (HDACs) even after MPP8 is switched off [13].

Our studies not only demonstrated MPP8 modulation by protein degradation, but also subcellular relocalization during HAdV infection (Fig. 4A). Immunofluorescence studies show that MPP8 localized to PML track-like structures (Fig. 4A, B) dependent on E1B-55K expression (Fig. 4C). In previous studies, E1B-55K was identified as viral factor counteracting several host factors, such as p53 [33]. In addition to proteasomal degradation, E1B-55K suppresses the transcription of p53-responsive genes by SUMOylation of p53. In this case, the function of E1B-55K is dependent on its interaction with PML-IV and PML-V [53]. This mechanism may be applicable to MPP8 regulation by E1B-55K since the immunoprecipitation assays indicate localization and interaction of MPP8 to PML tracks and E1B-55K mediated by PML-IV (Fig. 4D). In the context of HSV-1 infection genomes are placed at PML-NBs, so called vDCP-NBs [14]. The HUSH complex silences gene expression in those vDCP-NBs to promote latent HSV-1 infection. The counteraction of the HUSH complex is necessary to convert HSV- 1 from a latent state to active gene expression [43]. Similar to HSV-1 and other DNA viruses, HAdV place their genomes juxtaposed to the PML tracks [45]. In our studies, we observed MPP8 localization at PML tracks and simultanously at HAdV RCs (Fig. 4A, B). We hypothesize that MPP8 is present in two different fractions after HAdV infection. A fraction that is PML associated and the fraction found at the outer rim of RCs. We suggest that this is due to MPP8 that has already been recruited to the PML tracks via PML-IV and E1B-55K to inhibit the function of MPP8 and prepare it for subsequent proteasomal degradation (Fig. 1B, 2A, 2C, 4), whilst the remaining MPP8 associated with HAdV RCs could act as a negative factor mediating the deposition of H3K9me3 on the viral genome. This would be supported by the fact that we still see an effect on the viral progeny after MPP8 has been knocked down, implying that MPP8 cannot be completely counteracted during HAdV infection at earlier stages of infection (Fig. 3A). Overall, during HAdV infection, MPP8 localizes to PML tracks and viral RCs, likely in a manner dependent on E1B-55K and PML-IV. This interaction may represent a mechanism by which HAdVs suppress MPP8 function, similar to the known repression of p53 [46, 53]. Furthermore, these findings could contribute to the unravelling the mechanism of persistent/pseudo-latent HAdV infection, which is still poorly understood [43].

Taken together, this study investigates the role of the HUSH complex and specifically MPP8 the course of HAdV infection. This research demonstrates that MPP8 might promote heterochromatin formation on the viral genome, thus inhibiting viral replication and gene expression. Our finding show that MPP8 depletion in knock-down cells increases viral replication and transcription, most presumably involving an interplay between MPP8 and TASOR. Interestingly, the E1B-55K and E4orf6 viral proteins MPP8 and TASOR for proteasomal degradation via an E3 ubiquitin ligase, thereby supporting viral replication and gene expression. We provide evidence that MPP8 is recruited to PML tracks during infection, in dependency on viral E1B-55K expression. Additionally, we find MPP8 fractions at viral RC and suggest that there are two MPP8 fraction with unequal functions during HAdV infection.

## Material and Methods

### Cell culture

HEK293 (ECACC European Collection of Authenticated Cell Cultures; Sigma-Aldrich, No. 85120602-1VL), HEK293T (ATCC, No. CRL-3216), H1299 (ATCC, No. CRL-5803; [54]), H1299 shCRL, H1299 shMPP8 and HepaRG (Thermo Scientific; [55]) cells were maintained as a monolayer with Dulbecco’s Modified Eagle Medium (DMEM; Sigma-Aldrich) supplemented with 100 U of penicillin, 100 µg of streptomycin per ml (Thermo Scientific), 5% FCS (Thermo Scientific) for HEK293, HEK293T and H1299 cells, and 10% FCS for HepaRG in a 5% CO_2_ atmosphere at 37°C. For cultivation of HepaRG cells, medium was additionally supplemented with 5 µg/µl of bovine insulin and 0.5 µM hydrocortisone. All cell lines were frequently tested for mycoplasma contamination.

For generation of stable knockdown cell lines, HEK293T cells co-transfected with lentivirus packaging plasmids pLP1, pLP2, and pVSVg. In addition cells were wither transfected with pAPM-D4 miR30-L1221 (control shRNA; gift from Jeremy Luban; Addgene plasmid #115846; http://n2t.net/addgene:115846 ; RRID:Addgene_115846) or pAPM-D4 miR30-MPHOSPH8 ts2 (MPP8 knockdown; gift from Prof. Jeremy Luban, NY, USA (Addgene plasmid #115875; http://n2t.net/addgene:115875 ; RRID:Addgene_115875) [36]. To isolate the generated lentivirus particles, supernatant from transfected HEK293T cells were sterile filtered (0.45 µm) 4 days post transfection. H1299 cells were infected with lentiviral particles and 5 mg/ml polybrene. After 5 days, media was exchanged by selection media containing 5 µg/ml puromycin.

For inhibition of the proteasome, cells were treated with 25 µM MG-132 (Sigma-Aldrich) 16 h prior to harvesting.

### Human Adenoviruses

In this study, cells were infected with H5*pg*4100 wild-type (wt) virus [31], and mutant viruses depleted for distinct early viral gene products, H5*pm*4149 (ΔE1B-55K), H5*pm*4154 (ΔE4orf6), and H5*pm*4150 (ΔE4orf3) [56] at a MOI (multiplicity of infection) of 50 [57]. All viruses were provided by Prof. Thomas Dobner, LIV Hamburg. We propagated and titrated in HEK293 cells as previously published [57, 58]. Quantification of viral progeny was performed by quantitative immunofluorescence staining of the viral DNA binding protein E2A as described in [44, 58].

### Transient transfection of plasmid DNA

Subconfluent H1299 cells were transfected with a mixture of linear polythylenimine (PEI; 25 kDa and the respective DNA as described previously [31]. Plasmid DNA used in this study: pcDNA3.1, pLKO.1, pcDNA-p53 (kindly provided by Prof. Hans Will, UKE Hamburg), pLKO- FLAG-PML-I, pLKO-FLAG-PML-II, pLKO-FLAG-PML-III, pLKO-FLAG-PML-IV, pLKO-FLAG-PML-V, and pLKO-FLAG-PML-VI [59].

### Protein lysates, immunoprecipitation, and immunoblotting

For protein lysate preparation, cells were resuspended in an adequate volume of RIPA buffer (50 mM Tris-HCl [pH 8.0], 150 mM NaCl, 5 mM EDTA, 1% [vol/vol] Nonident p-40, 0.1% [wt/vol] SDS, and 0.5% [wt/vol] sodium deoxycholate) supplemented with protease inhibitors 0.2 mM PMSF, 1 mg/mL pepstatin A, 5 mg/mL aprotinin, and 20 mg/mL leupeptin. Samples were vortexed every 10 min and incubated on ice for a total of 30 min. After incubation, the samples were subjected to sonication in three cycles (*Diagenode* Bioruptor Plus, high setting 30 s on and 30 s off) at 4°C. Cell debris was pelleted for 3min at 11,000 rpm and 4°C. For cell lysates the required amount of protein were boiled for 3 min at 95°C in laemmli buffer and subjected to immunoblotting as previously described [60].

For immunoprecipitation, 2000 µg of protein lysate and 50 µl of SureBeads^TM^ (Biorad) per sample were used. First, magnetic SureBeads^TM^ were coupled with 5 µl of mouse mAb MPP8 (sc-398598; Santa Cruz Biotechnology) for 30 min at RT. Antibody-coupled magnetic beads were added to the lysate samples and rotated for 1 h at RT. After incubation samples were washed thrice with PBS-T, boiled for 5 min at 95°C in 2x laemmli buffer and subsequently subjected to immunoblotting as previously described [60].

Primary antibodies used for immunoblotting are mouse mAb AC-15 (anti-β-actin; Sigma-Aldrich), mouse mAb MPP8 (sc-398598; Santa Cruz Biotechnology), rabbit pAb MPP8 (HPA040035; Sigma-Aldrich), rabbit pAb PPHLN1 (HPA0038902; Sigma-Aldrich), rabbit pAb TASOR (FAM208A; NBP1-90673; Novus Biologicals), rabbit pAb NP220 (A301-547A; Biomol) mouse mAb Ubiquitin (P4D1; Cell Signaling), rabbit pAb PML (ab72137, abcam), mouse mAb FLAG(M2) (F1804, Sigma Aldrich), mouse mAb E1A (M73; sc-25, Santa Cruz Biotechnology; [61]), mouse mAb E1B-55K (4E8, [62]), mouse mAb E2A (B6-8, [63]), rat mAb E4orf3 (6A11, [64]), mouse mAb E4orf6 (RSA3, [65]), and rabbit pAb Capsid (L133, [66]). Secondary IgG antibodies (Jackson/Dianova) conjugated to horseradish peroxidase (HRP) were diluted 1:10000 in 3% milk powder in PBS-T.

### Quantitative real-time PCR analysis

Subconfluent H1299 shCRL and shMPP8 cells were infected with wild-type virus and harvested according to the experimental setup. Total RNA isolation was performed with TRIzol reagent (Thermo Scientific) according to the manufacturer’s protocol. For reverse transcription of the isolated RNA, 1 µg of RNA was submitted to the *Promega Reverse Transcription System* according to the manufacturer’s protocol.

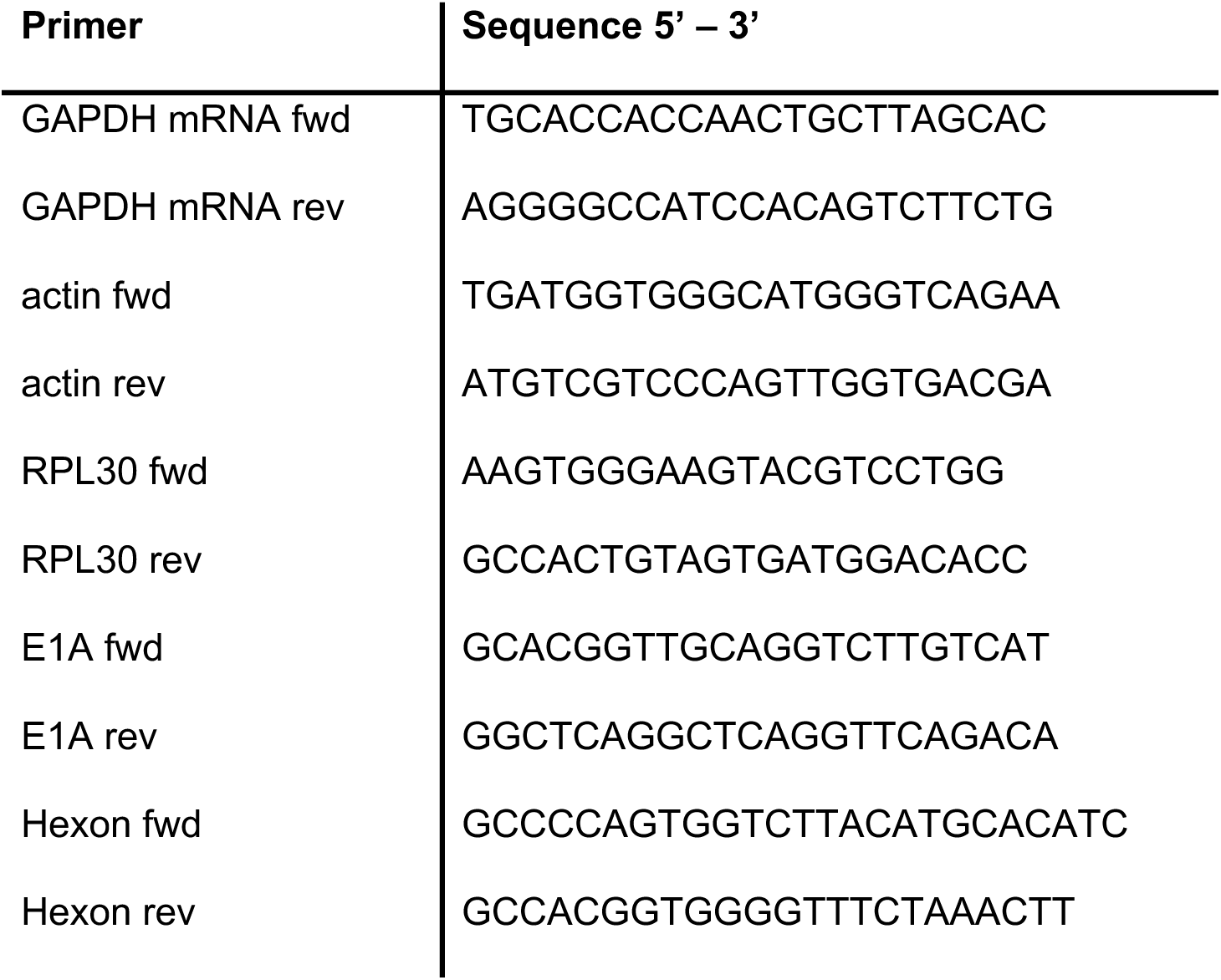

Quantitative RT-PCR was performed in a 96well plate (Biorad) with a first-strand method. As a template 5 µl of a 1/10 cDNA dilution (mRNA) were mixed with 10 pmol/µl of each primer and 5 µl of the Luna Universal Master Mix (NEB) per sample. The PCR conditions were set as follows: 2 min at 95°C and 40 cycles of 15 s at 95°C, and 45 s at 60°C. Viral E1A and Hexon transcript levels were calculated in relation to the cellular housekeeping transcripts actin, GAPDH, and RPL30 with the ΔΔCt method.

### Indirect immunofluorescence

To visualize protein localization, indirect immunofluorescence assays were performed as described previously [44]. In brief, cells were seeded on glass coverslips and treated according to the experimental setup. At the indicated time points cells were fixed with 4% paraformaldehyde in PBS, permeabilized in 0.5% Triton-X100 (Sigma-Aldrich). Subsequently, coverslips were blocked in TBS-BG and immunostained with the respective antibodies E4orf3 rat mAb 6A-11 [66], E2A rabbit mAb (kindly provided by Prof. Ron T. Hay, Dundee, Scotland), and MPP8 mouse mAb (sc-398598; Santa Cruz Biotechnology). Secondary antibodies conjugated with Alexa647 anti-rat, Alexa568 anti-rabbit, and Alexa488 anti-mouse were purchased from Dianova and Invitrogen, respectively. DAPI (Sigma-Aldrich) was used for nuclear staining and coverslips were mounted in Mowiol 4-88 (Carl Roth). Fluorescence images were taken at a *Zeiss* LSM 900 Airyscan 2. Images were processed analyzed using the *Zeiss* ZEN 3.0 blue edition software.

## Acknowledgement

This work was funded by the Deutsche Forschungsgemeinschaft (DFG, German Research Foundation) in the framework of the Research Unit FOR5200 DEEP-DV (443644894) project SCHR 1479/5-1 to S.S. and and STA357/8-1 to T.S..

## Author Contribution

Conceptualization: SS, JM, MK; Methodology: JM, MK, AK, TG; Investigation: JM, MK, AK, TG; Formal analysis: SS, JM, MK; Resources: AG, TS; Data Curation: SS, JM, MK; Writing Original Draft: MK, JM, SS; Writing – Reviewing & Editing: JM, SS; Visualization: MK, JM, SS; Project administration: SS; Supervision: SS; Funding Acquisition: SS

